# Where do the ligands bind? Co-folding bitter taste GPCRs with the BitterDB chemical space

**DOI:** 10.64898/2026.06.26.734512

**Authors:** Evgenii Ziaikin, Masha Y. Niv

## Abstract

Bitterness is a key taste modality mediated in vertebrates by TAS2R G-protein-coupled receptors, which also function in diverse extraoral tissues. Recent cryo-EM structures have revealed a non-classical intracellular pocket in TAS2R14, raising the question of whether ligand pocket choice can be predicted computationally and what sequence features control it. Here we evaluate the Boltz-2 co-folding framework on all currently available agonist-TAS2R cryo-EM complexes and show that it correctly identifies the experimentally observed binding pocket for 12 of 15 pairs, including intracellular binding that docking into predicted receptor models fails to reproduce. Focusing on aristolochic acid, which binds intracellularly to TAS2R14 and extracellularly to TAS2R43, we use a series of in silico morphing experiments to pinpoint transmembrane helices 3 and 7, and specific residues within them, as key determinants of pocket preference. Extending the analysis to ∼1,500 agonist-receptor associations from BitterDB, we find that while most receptors are predicted to bind agonists predominantly in the extracellular pocket, several TAS2Rs may have both extracellularly and intracellularly binding ligands. Finally, by fine-tuning the Boltz-2 affinity module on ∼7,000 positive and negative experimental data points, we obtain a TAS2R-specific classifier that improves AUROC from 0.54 to 0.82 and average precision from 0.24 to 0.58 on a validation set.

## Introduction

Bitterness is one of the basic taste modalities that determine food preferences^1^. It is mediated via TAS2R receptors, which are part of a large superfamily of transmembrane G-protein coupled receptors (GPCRs)^2^. There are 26 variants of TAS2Rs in the human body^3,4^, each with particular repertoire of ligands^5^ and individual patterns of expression in tissues other than the oral cavity^6^, suggesting that this family may have other biological roles besides taste perception^7–9^.

Following advances in cryo-EM structural biology for GPCRs^10^, there is currently an increasing number also of TAS2R cryo-EM structures^11^. The first cryo-EM structure solved among TAS2Rs was the strychnine complex with TAS2R46^12^, which confirmed the canonical binding pocket for all GPCRs in the canonical (extracellular) region of the receptor. The same situation was observed for WWW tripeptide complex with TAS2R4^13^, salicin complexes with TAS2R16^14^, aristolochic acid complexes with TAS2R43^15^, and other ligands with TAS2R46^16^. However, TAS2R14 solved complexes with cholesterol^17–19^, compound 28.1^17–19^, flufenamic acid^18,20^, aristolochic acid^18^, and others^16^, revealed binding in an extracellular or intracellular pocket. Moreover, solved complexes for flufenamic acid^20^ and a related compound 28.1^19^ with TAS2R14 demonstrated two copies of the same ligand binding in two pockets simultaneously. This accumulating data raises the question about the determinants of the location of the ligand binding in TAS2Rs, and the ability to predict where the ligands bind.

The most up-to-date information on bitter TAS2R receptors and their activating ligands is collected in BitterDB^21^, a publicly available database that combines information from scientific literature. BitterDB currently contains more than 2,200 unique compounds, over 1,700 experimentally confirmed bitterant–receptor functional assays and includes above 60 different species of organisms besides humans.

Despite numerous successful structure-based screening campaigns that relied on predicted three-dimensional structures of TAS2Rs^20,22–29^, the intracellular binding pocket itself remained unexplored until its identification in recent cryo-EM structures^17–20,30^. Subsequent cryo-EM^17–20,30^ and mutagenesis studies^26,31,32^ have now established that this pocket plays a key role in the activation of TAS2R14 by certain ligands.

The aim of this work is to estimate where the ligands bind to TAS2Rs, and which regions are particularly important for determining the binding location. We chose Boltz-2^33^, as a representative, easy to use co-folding method that allows for relatively quick and accurate prediction of protein-ligand complexes^34^. We first validated its performance in recapturing experimental locations against the available сryo-EM structures of TAS2Rs. Next, we used computational morphing and co-folding to identify key positions dictating extracellular or intracellular binding for a specific case study and mapped the bitter chemical space to TAS2Rs extracellular or intracellular binding sites. Finally, we fine-tuned the affinity module in Boltz-2 to distinguish true positives (agonists) from true negatives (non-agonists).

## Results

### Reproduction of experimental poses for the TAS2R family using co-folding

Prior to the publication of cryo-EM structures of TAS2s, we^20,25–29^ and others^22–24^ have used various modelling approaches (MEDELLER^35^, I-TASSER^36^, AlphaFold^37^) for docking of ligands to TAS2R14 model. Interestingly, docking of molecules to a computationally predicted model of TAS2R14, does not reproduce the intracellular binding of ligands. This is also the situation even when docking to receptor structures predicted by the modern protein folding tools such as AlphaFold3^38^ and Boltz-2^33^, result in binding poses within 7.0 to 13.5 RMSD from the experimental ones (Supplementary Figure 1).

We asked whether co-folding will perform better. To date (June 2026), a total of 37 cryo-EM structures have been solved for TAS2Rs (33 available and 4 “on-hold”), which form a set of 15 agonist-TAS2R pairs (Supplementary Table 1,2). New structures from a recent preprint^30^ presented 4 experimental cryo-EM structures for TAS2R14, including two with new ligands cholesteryl hemisuccinate in the extracellular pocket and diiodosalicylic acid in the intracellular. The structures are still unavailable, but pockets were described in the preprint^30^. Thus, 15 (2 from the preprint^30^) ligand-protein pairs were predicted using Boltz-2^33^, which was trained on the 2023 PDB dataset that only included 3 TAS2R46’s cryo-EM structures with strychnine^12^ (Figure 1). Root Mean Square Deviation (RMSD) was used to evaluate the quality of the predicted pose.

**Figure 1.**
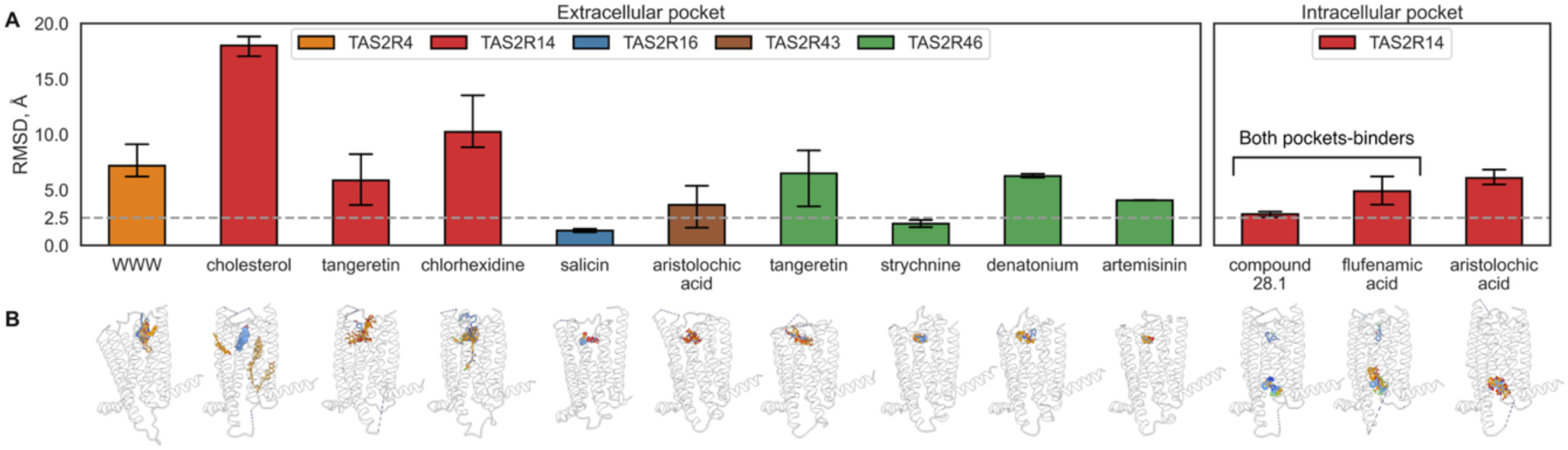
(A) RMSD values between ligands in co-folded agonist-TAS2R complexes and their experimental cryo-EM references for the extracellular and the intracellular pocket. The gray dashed line represents a 2.5 Å threshold indicating the cutoff for well-reproduced poses. (B) Aligned experimental (blue) and predicted (orange) binding poses of TAS2R agonists.

As seen in Figure 1, Boltz-2 reproduced the binding pockets for all TAS2R ligands studied, except for cholesterol and chlorhexidine. For the diiodosalicylic acid-TAS2R14 complex, Boltz-2 correctly predicts binding within the intracellular pocket, while for cholesteryl hemisuccinate (a cholesterol analog), the ligand is positioned incorrectly (Supplementary Figure 2).

For cases where ligands can bind to both binding pockets, namely TAS2R14 with compound 28.1 and flufenamic acid, Boltz-2 places the ligands only in the intracellular pocket (Figure 1B), with RMSD of 2.8 ± 0.4 Å and 4.9 ± 1.8 Å respectively. This preference for the intracellular pocket is consistent with mutagenesis results, which supported the dominant role of the intracellular pocket in the activation of TAS2R14 by these agonists^20,32^. The aristolochic acid bound in several locations of the intracellular region of TAS2R14^18,30^. Boltz-2 reproduces a particular intracellular pose (called pose 1), which is found in all solved cryo-EM complexes of aristolochic acid with TAS2R14 (Figure 1B).

### Morphing of TAS2R14 and TAS2R43 binders of aristolochic acid

Co-folding of TAS2Rs with aristolochic acid (AA) reveals the correct ligand location both for intracellular binding in TAS2R14, and extracellular binding in TAS2R43(Figure 1). What are the sequence signals that direct the (correct) prediction to either extracellular or intracellular pocket? We performed a series of mutual transformations of the TAS2R14 and TAS2R43 pair to identify the sequence changes that lead to change in predicted ligand placement. Few morphing strategies were used: a) replacement of the extracellular or the intracellular pockets, b) replacement of individual TMs, and c) step-by-step morphing of individual TMs affecting the AA binding mode (Figure 2 A-C).

**Figure 2.**
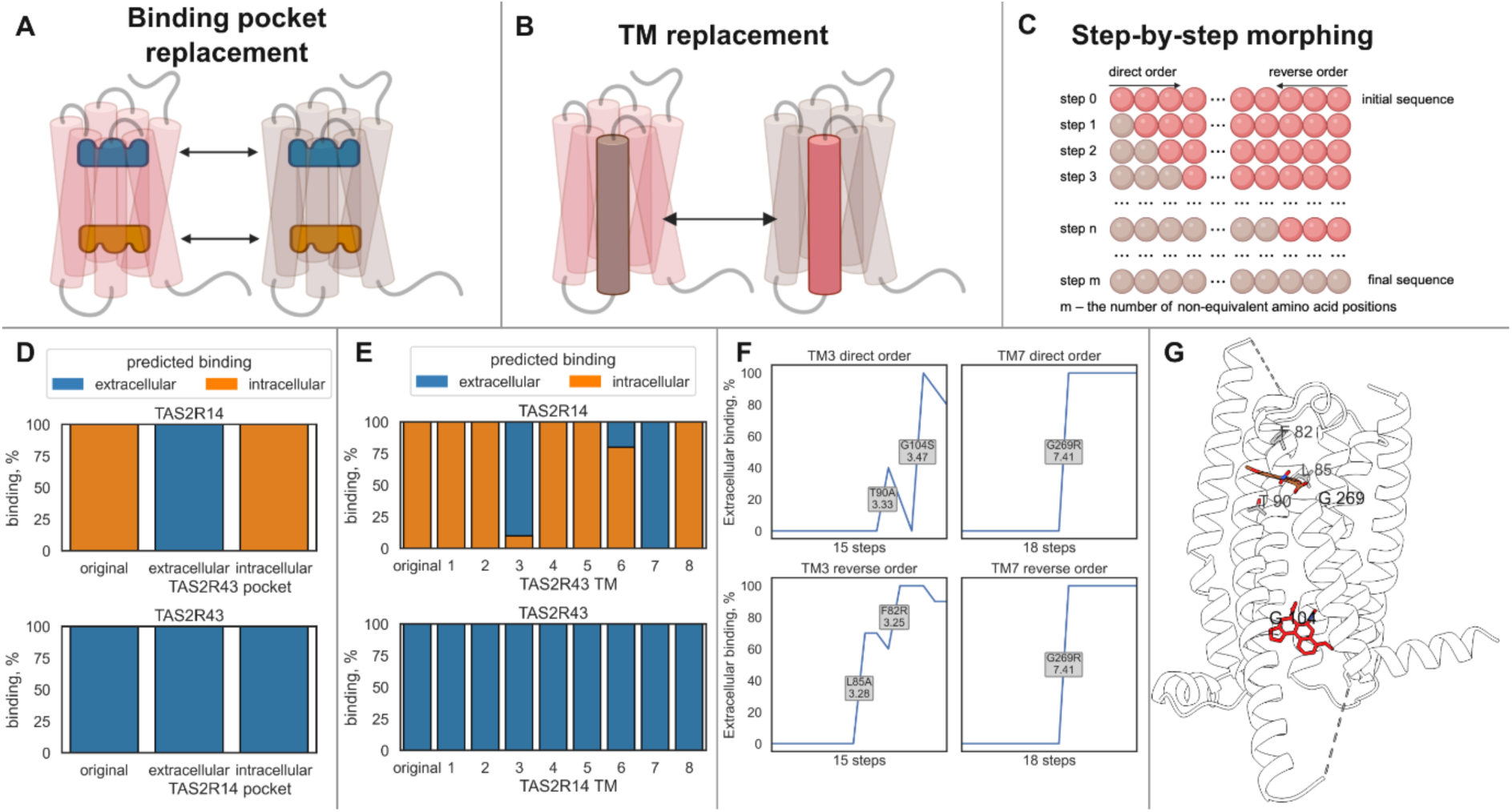
Visualization of the morphing scenarios performed for TAS2R14 and TAS2R43 with aristolochic acid (AA): (A) binding pocket replacement, (B) TM replacement, (C) step-by-step morphing. Results of morphing scenarios predictions: (D) binding pocket replacement, (E) TM replacement, (F) step-by-step morphing of TM3 and TM7 of TAS2R14 into TAS2R43 corresponding TMs. The y-axis shows the percentage of binding among the 10 predicted samples in the extracellular pocket (blue) and in the intracellular pocket (orange). Mutations that lead to a significant increase in extracellular AA binding in the TAS2R14 chimeric protein are shown. (G) Location of residues in TAS2R14 whose mutations lead to an increase in the predicted extracellular binding of AA. The experimental poses of AA in TAS2R14 (8XQO) are shown in red, and those in TAS2R43 (9OXA) are shown in brown.

Replacing the extracellular pocket of TAS2R14 with that of TAS2R43 completely shifts AA binding to the extracellular region, whereas replacing the intracellular pocket of TAS2R14 with that of TAS2R43 does not alter AA binding in the intracellular region. In the case of TAS2R43, none of the replaced binding pockets affected AA binding in the extracellular pocket (Figure 2D). This may suggest that TAS2R14 binds AA in the intracellular pocket not because its intracellular pocket is better for AA than the intracellular pocket of TAS2R43, but because TAS2R14 extracellular pocket is less suitable for it.

When individual TMs in TAS2R14 were replaced with TMs from TAS2R43, a complete shift in the AA binding mode from intracellular to extracellular was observed for TM3 and TM7, and a partial shift was also observed for TM6 (Figure 2E). In the case of substitutions in TAS2R43, none of the substituted TMs affected the AA binding mode.

Next, a step-by-step cumulative replacement was performed on each of these TMs, starting from either N or C terminus, and co-folding results predicted mutations that led to extracellular AA binding in the TAS2R14 chimeric protein (Figure 2F). The aligned sequences of TM3 and TM7 involved in the step-by-step morphing are shown in Supplementary Figure 3.

We did not find mutations in the binding site or individual TMs, that were predicted to revert TAS2R43 to an intracellular site AA binder. Conversely, several combined mutations or replacement of either TM3 or TM7 in TAS2R14 with the analogous parts of TAS2R43, can lead to switch of AA binding to the extracellular site.

Among the determining positions in TAS2R14 are F82^3.25^, L85^3.28^, T90^3.33^ and G269^7.41^ reside in the extracellular pocket, while G104^3.47^ is located in the intracellular pocket (Figure 2G). In TAS2R43, these residues are R81^3.25^, A84^3.28^, A89^3.33^, R268^7.41^, and S103^3.47^. The G104A^3.47^ mutation is known experimentally to impair AA binding in the intracellular pocket^18^, and the G269R^7.41^ substitution likely enhances extracellular binding by creating a residue that directly contacts AA in TAS2R43^15^. The mutations F82R^3.25^, L85A^3.28^, and T90A^3.33^ remodel the extracellular pocket of TAS2R14 to more closely resemble TAS2R43’s, though these positions have previously been studied without producing changes in AA activation levels^26^.

### Predicted intracellular pockets in other TAS2Rs

The overall successful identification of intracellular ligand binding without additional constraints during co-folding highlighted the potential of Boltz-2 for predicting new candidates of intracellular ligands for other TAS2Rs.

We used Boltz-2 to predict 5 complexes for each of the experimentally established 1,555 agonist-TAS2R associations in BitterDB^21,33^, which corresponded to 139 TAS2Rs from several species, that are associated with at least one ligand. To ensure greater prediction reliability, we used the interface predicted TM score (ipTM), a metric that indicates the accuracy of interfaces in the predicted structure and is evaluated using a confidence module. In the original Boltz-2 paper, an ipTM threshold below 0.75 was used to flag unreliable predictions, and we use it in the same way in this work. As a result, we retained 1,511 agonist-receptor pairs, with 5 Boltz-2 predicted complexes each. The percentage of intracellular binding was calculated for each TAS2R using Boltz-2 predictions (Figure 3).

**Figure 3.**
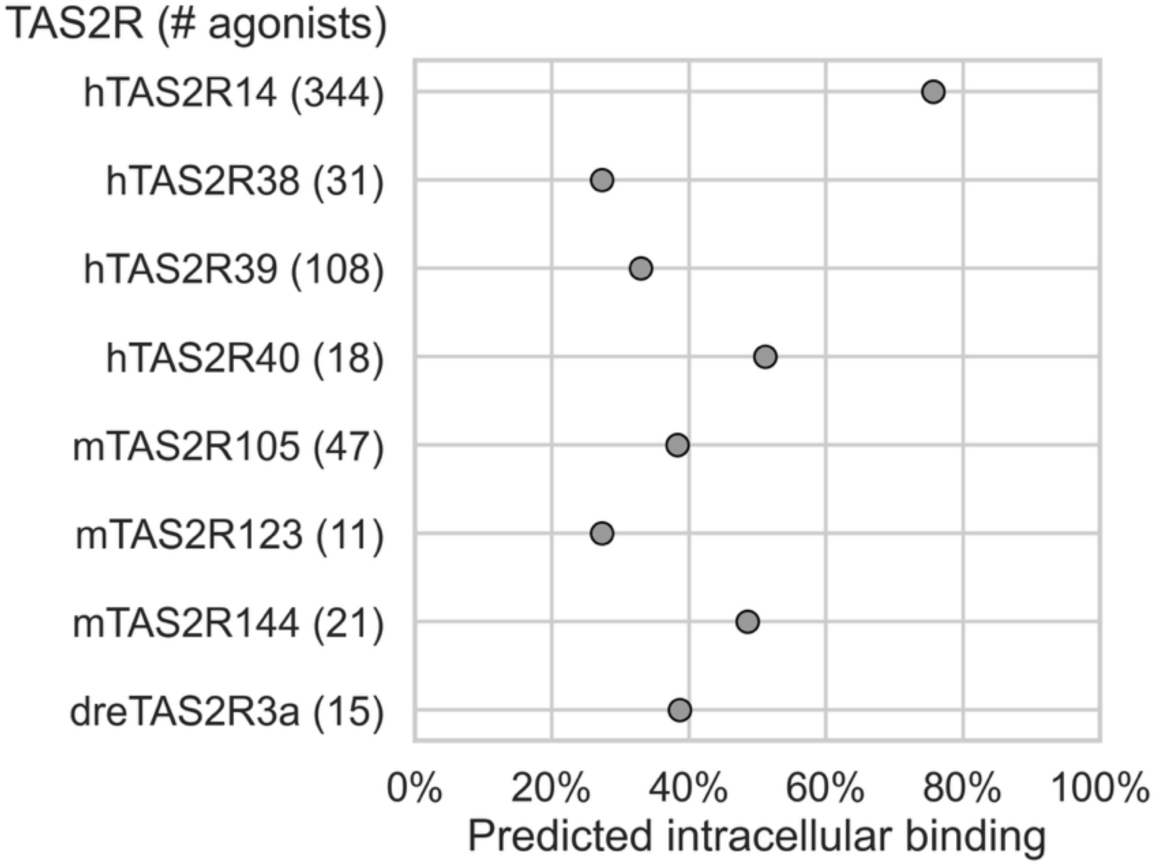
Distribution of predicted intracellular binding of TAS2R agonists. The number of agonists is indicated in brackets after the receptor name. Only those TAS2Rs for which more than 10 agonists are known, and the percentage of intracellular binding exceeds 20% are shown. Prefixes indicate species: h – Human (Homo sapiens), m – Mouse (Mus musculus), dre – Fish (Danio rerio).

For most TAS2Rs (118 out of 139), Boltz-2 predicts canonical extracellular binding, (Figure 3A). Among human TAS2Rs, TAS2R38, TAS2R39, and TAS2R40 stand out as a group with intracellular fractions of 27–51%. The aligned sequences of the extracellular and intracellular pockets of human TAS2Rs are shown in Supplementary Figure 4. Among the new candidates for binding within the intracellular pocket are several TAS2R receptors from the mouse (*Mus musculus*, prefix m) and zebrafish (*Danio rerio*, prefix dre) (Figure 3A), suggesting that the propensity for intracellular binding is not limited to human receptors.

### Affinity values for TAS2R agonists and non-agonists

We next asked whether the method could provide informative results for ligands whose association with TAS2Rs has not been experimentally established. Therefore, in addition to applying co-folding and affinity prediction with Boltz-2 to the dataset of 1,555 agonist-TAS2R pairs, we also analyzed a dataset of 6,924 non-agonist-TAS2R pairs (Figure 4).”. Although the authors of Boltz-2 note in the original repository (https://github.com/jwohlwend/boltz) that the predicted affinity module values are intended only to assess the efficiency of active compounds, we were expecting some bias in the estimates for TAS2R agonists vs. non-agonists. However, for all human TAS2Rs (Figure 4A), the estimates for agonists and non-agonists were in similar ranges (Figure 4B).

**Figure 4.**
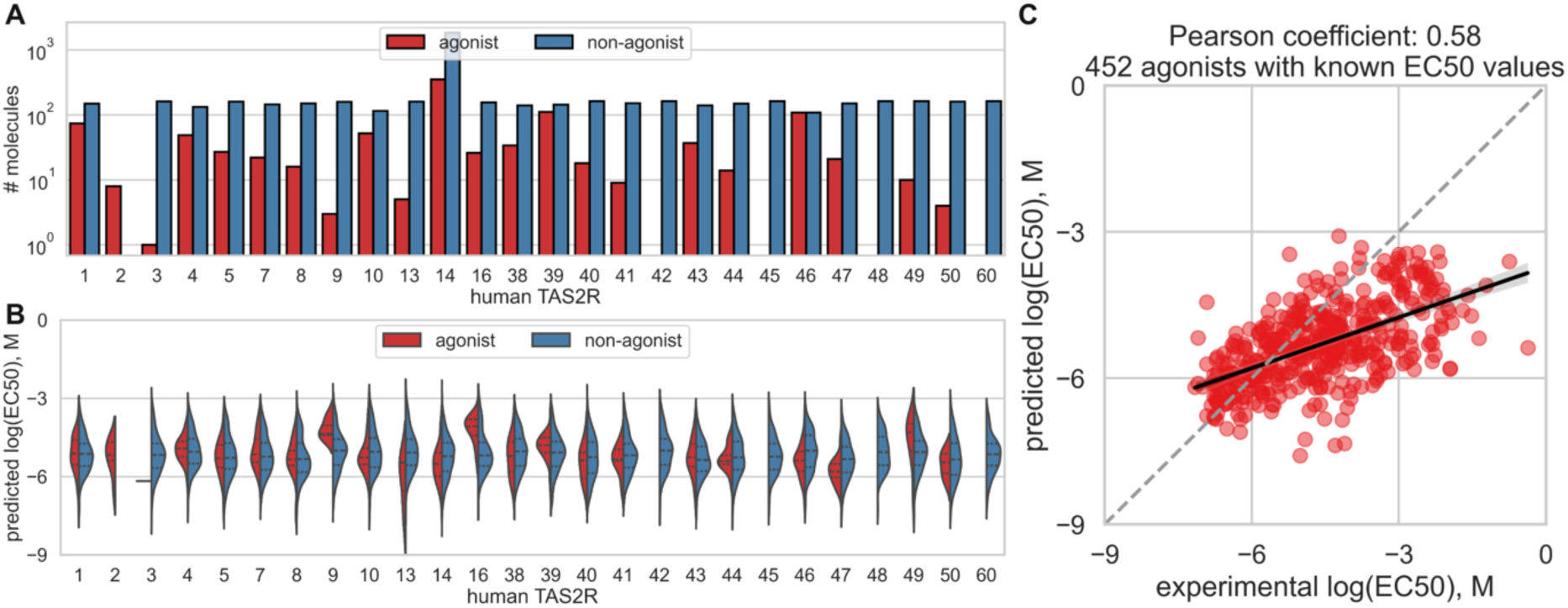
(A) Numbers of agonists and non-agonists of human TAS2Rs taken from articles cited in BitterDB^21^. (B) Predicted affinity values for agonists and non-agonists of human TAS2Rs. (C) Correlation between predicted and experimental TAS2R affinity values for agonists.

For 452 agonist-TAS2R pairs, the affinity module predictions show a Pearson coefficient (Pearson R) of 0.58 with experimental value (Figure 4C). This is somewhat lower than for the GPCR targets studied in the original work (Pearson R = 0.70–0.73)^33^ and other studies (Pearson R = 0.98)^39^, which used a smaller number of experimental data points. The accuracy of the predicted affinity values for individual TAS2Rs varies (Supplementary Figure 5), and typically did not improve for TAS2R with many known ligands. But no direct correlation was observed between the number of agonists with known EC50s and Pearson R.

### Fine-tuning Boltz-2 affinity module

We next used this set of TAS2R agonists and non-agonists to fine-tune Boltz-2 affinitiy module. The set used 1,557 positive and 6,924 negative TAS2Rs associations. The data was split into train, and validation sets in an 80:20 proportion, keeping the original distribution of positive and negative examples for each receptor in each of the subsets. The results were calculated for the validation set. The training curves are shown in Supplementary Figure 6. The metrics for the best model are shown in Supplementary Table 3. A significant improvement was achieved (Figure 5), reaching ROCAUC = 0.82 (originally 0.54) and AP = 0.58 (originally 0.24). Prediction accuracy increased for all receptors (Supplementary Figure 7).

**Figure 5.**
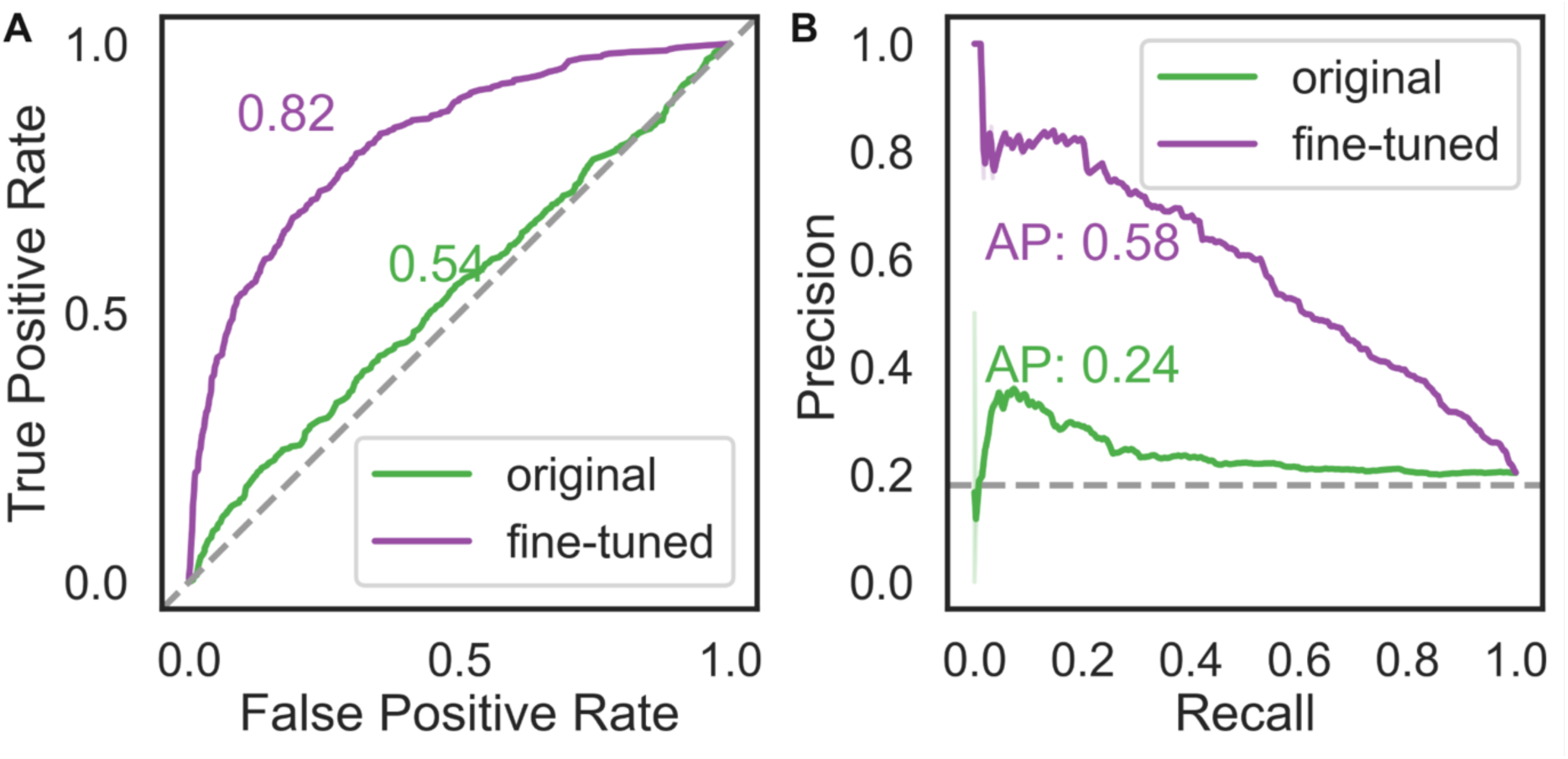
Results of fine-tuning the affinity module: (A) ROC curve and (B) precision-recall curve for the original and fine-tuned models. The dashed lines indicate the performance of a random model; for the PR curve, this was calculated as the proportion of the positive class in the original dataset, which was 0.18.

## Discussion

Boltz-2 was evaluated for its accuracy in reproducing experimentally validated binding poses for agonists of the TAS2R bitter receptor family. For 12 of 15 agonist-TAS2R pairs with solved experimental structures, Boltz-2 correctly identified the binding pocket, in some cases with RMSD values below 2.5 Å – a strong performance given that only TAS2R46 cryo-EM structures were included in its training set.^33^. Notably, Boltz-2 correctly identified the intracellular binding pocket for TAS2R14 agonists, a rare feature among GPCRs and one that was missed by classical docking into computationally predicted receptor models, and may be used in the future for identifying potential intracellular, including endogenous, ligands.

Three complexes that were predicted incorrectly, involved TAS2R14 with cholesterol, chlorhexidine, or cholesteryl hemisuccinate ligands. Cholesterol is routinely used to stabilize GPCRs during cryo-EM sample preparation, localizing on the outer transmembrane surface in interfacial grooves^40,41^, and is found complexed in a wide variety of GPCR structures^42^. These were also the locations predicted for TAS2R14 with cholesterol and its derivatve cholesteryl hemisuccinate, but cryo-EM structures showed them in the canonical extracellular pocket of TAS2R14. The large error in reproducing the chlorhexidine may relate its high conformational flexibility

In the cryo-EM structures, aristolochic acid (AA) binds extracellularly in TAS2R43 but intracellularly in TAS2R14, and Boltz-2 correctly recovered these distinct modes, enabling morphing-based dissection of sequence determinants of pocket preference. Swapping pockets in silico showed that replacing the extracellular, but not the intracellular, pocket of TAS2R14 with that of TAS2R43 redirected AA to the extracellular site, indicating that TAS2R14 intracellular binding may arise from an unfavorable extracellular environment rather than a uniquely favorable intracellular pocket, consistent with AA ∼2-fold lower EC50 at TAS2R14. At the helix level, substituting TM3 or TM7 of TAS2R14 with their TAS2R43 counterparts likewise shifted AA toward extracellular binding, whereas no single TM swap in TAS2R43 altered its extracellular preference; although Boltz-2 is relatively insensitive to single-residue changes, cumulative TM3/TM7 replacements nevertheless highlighted positions whose mutation markedly increased extracellular AA-binding predictions in the TAS2R14 background.

The ability of Boltz-2 to recover intracellular ligand binding without explicit geometric constraints suggested it could be used to screen TAS2Rs more broadly. Applied to 1,511 high-confidence agonist-TAS2R pairs from BitterDB, Boltz-2 assigned most receptors to extracellular binding, but for several, including TAS2R14, TAS2R38, TAS2R39 and TAS2R40, a substantial fraction of agonists was predicted to bind intracellularly (Figure 3). Finally, although the original Boltz-2 affinity module was not optimized for binary agonist classification, fine-tuning its classification head on the 6,783 TAS2R associations (train set) dataset from BitterDB markedly improved performance, indicating that this architecture can be effectively adapted to a specific GPCR family, even when training on functional assay data, as is the case for TAS2Rs.

In 2025, approximately 36% of approved drugs targeted GPCRs^43^, underscoring the long-standing pharmaceutical relevance of this receptor superfamily. The discovery of a functional intracellular binding pocket in at least one TAS2R, and its probable presence in additional family members, opens new avenues for modulation. Further elucidation of the structural determinants governing intracellular pocket formation and their coupling to receptor activation may reveal general principles that extend beyond TAS2Rs and inform GPCR biology and drug discovery more broadly. Notably, TAS2Rs themselves are emerging as therapeutically relevant targets^44–46^, and their relatively compact architecture, well-characterized ligand repertoires, and rapidly expanding cryo-EM structural dataset make them an especially intriguing and timely target.

## Acknowledgements

We thank Prof. Dina Schneidman and Ben Shor for helpful ideas that sparked the initiation of this project.

## Methods

### Docking of intracellular TAS2R14 agonists to predicted TAS2R14 structure

The sequence of TAS2R14, smiles for compound 28.1, flufenamic acid, and aristolochic acid were obtained from BitterDB^21^. A 3D model of the TAS2R14 receptor was predicted using AlphaFold3^38^ and Boltz-2^33^ (June 2026). Ligand preparation was performed in LigPrep (Schrödinger Release 2026-2: LigPrep, Schrödinger, LLC, New York, NY, 2025), receptor preparation was performed using the Protein Preparation Workflow (Schrödinger Release 2026-2: Protein Preparation Workflow; Epik, Schrödinger, LLC, New York, NY, 2024; Impact, Schrödinger, LLC, New York, NY; Prime, Schrödinger, LLC, New York, NY, 2025), and binding pockets were identified using SiteMap (Schrödinger Release 2026-2: SiteMap, Schrödinger, LLC, New York, NY, 2025). Docking was performed using Glide (Schrödinger Release 2026-2: Glide, Schrödinger, LLC, New York, NY, 2025) with standard settings.

### Dataset of TAS2Rs cryo-EM complexes

37 cryo-EM structures of TAS2R complexes were obtained from the PDB^47^. A complete list of the structures used is provided in Supplementary Table 1, 2.

### Dataset of TAS2R agonists and non-agonists

Data curation was performed according to the data preparation protocol described in the original Boltz-2 paper^33^. BitterDB^21^ was used to build a set of agonists and non-agonists. For agonists, SMILES, associations with TAS2Rs, protein sequence, and experimental EC50 values were extracted. To extract non-agonists, articles contained in BitterDB with experimentally confirmed negative associations with TAS2Rs were used. For SMILEs in both sets, standardization was performed using the ChEMBL Structure Pipeline^48^, followed by the removal of salts; when duplicates were found, records were merged, and compounds with more than 50 heavy atoms were discarded. If multiple EC50 values were available for an agonist-TAS2R pair, their average value was calculated. Next, the set of agonists and non-agonists was combined, and in the case of contradicting associations, preference was given to positive ones. The final set contained 1,555 positive agonist-TAS2R associations, of which 452 have known EC50 values, and 6,924 negative non-agonist-TAS2R associations.

### Co-folding of TAS2R protein-ligand complexes using Boltz-2

Currently (June 2026), a total of 37 (4 on hold) cryo-EM complexes of TAS2Rs with 32 containing a ligand (3 on hold). Based on this list, a set of 15 ligand-TAS2R pairs was prepared (Supplementary Table 2). For the co-folding of TAS2R protein-ligand complexes using Boltz-2^33^, the code and model weights provided by the official repository (https://github.com/jwohlwend/boltz) were used. Complex prediction and affinity value estimation were performed using default parameters, except for the diffusion samples, which were set to 5.

### Calculation of RMSD

Structure alignment, calculation of Root Mean Square Deviation (RMSD) and generation of ligand-protein interaction profiles were performed using MDAnalysis^49^, RDKit^50^, and PLIP^51^. When calculating the RMSD between the Boltz-2-predicted complexes and the experimental cryo-EM complexes, all available PDB entries for each ligand-TAS2R pair were used (Supplementary Table 1,2).

### Determination of the binding region

The classification of the ligand binding region into extracellular and intracellular was based on the proximity of the residues with which the ligand interacts to the extracellular and intracellular loops of the receptor. The residues corresponding to the loops for TAS2Rs were taken from the relevant entries in UniProt^52^.

### Morphing process of TAS2R-TAS2R

To perform the morphing process, the source sequence and target sequence of the corresponding TAS2Rs were used. These sequences were aligned, and then step-by-step morphing of non-equivalent positions in the MSA was performed using one of the following operations: replacement of the source amino acid with the target amino acid, deleting the source amino acid if there is a gap at the position instead of the target amino acid, and adding the target amino acid if there is a gap at the position instead of the source amino acid.

### Fine-tuning of the Boltz-2 affinity module

To fine-tune the affinity modules, we used a dataset of TAS2R agonists and non-agonists that had previously been co-folded using Boltz-2^33^. This resulted in a dataset of 1,555 positive associations and 6,924 negative associations (total 8,479 experimantal data points). The data was split into train and validation sets in an 80:20 proportion, keeping the original distribution of positive and negative examples for each receptor in each of the subsets. Thus, there were 6,783 associations in the train set and 1,696 associations in the validation set.

EC50 values for positive associations were not used in the fine-tuning of the affinity module due to the low density of such data in the final dataset. Only 452 of the 8,479 associations had experimental EC50 values, accounting for approximately 5% of the entire dataset. For this reason, fine-tuning was performed only on the classification head of the affinity module.

The Boltz-2 weights, provided with the code from the official repository (https://github.com/jwohlwend/boltz), were used as the starting model for fine-tuning the affinity module. Due to the small number of examples compared to the original work^33^, in which ∼4.5 million examples were used to train the affinity module, fine-tuning was performed only for affinity module ensemble member 2, which has less trainable parameters compared to ensemble member 1. Thus, the number of trainable parameters was ∼3.25 million.

The BCEWithLogitsLoss was used as the loss function with a positive weight parameter equal to 4.56, calculated as (1 – 0.18) / 0.18, where 0.18 is the proportion of positive examples in the dataset. Fine-tuning was performed over 20 epochs with a learning rate of 1e-4.

After constructing the path, a list of step-by-step morphed sequences was generated, for which co-folding was performed with standard settings, but with 10 diffusion samples. Based on the predicted complexes, dependencies of the target property (e.g., the share of ligand binding in the intracellular pocket) on the step in the morphing path were established.

## Supplementary Material

**Supplementary Figure 1.**
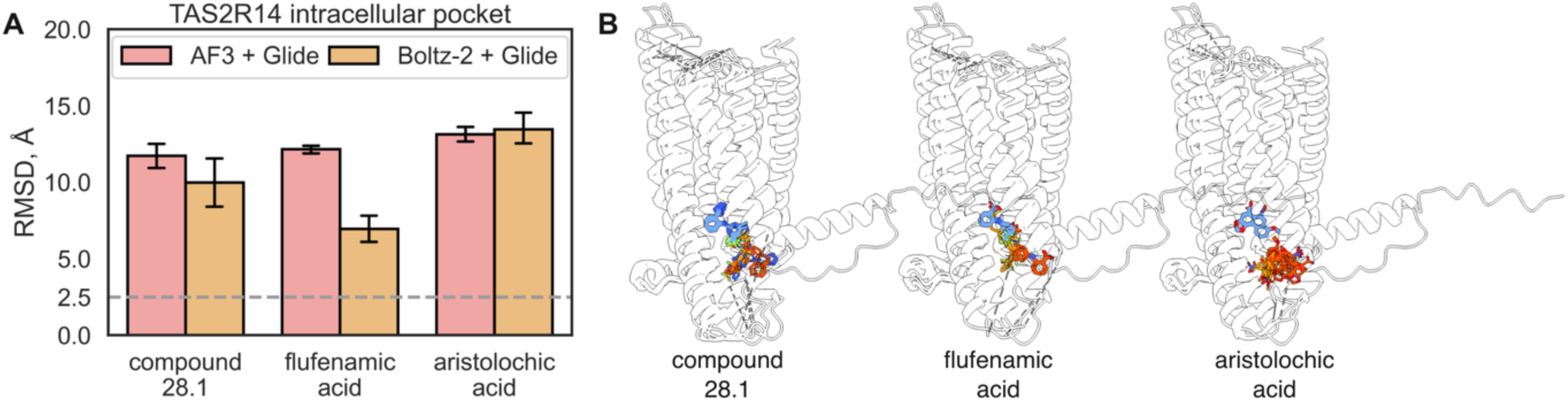
Accuracy of reproducing experimental intracellular conformations using AlphaFold3 (AF3) or Boltz-2 receptor folding followed by ligand docking with Glide for compound 28.1, flufenamic acid, and aristolochic acid in TAS2R14. (A) RMSD values against experimental cryo-EM structures. (B) Aligned experimental complexes (blue), folded receptor by AF3 and docked ligand by Glide (red), folded receptor by Boltz-2 and docked ligand by Glide (orange).

**Supplementary Table 1.** Unique “ligand-TAS2R” pairs with known experimental cryo-EM structures and results of binding site prediction using Boltz-2. Cases where Boltz-2 correctly predicted the binding pocket are highlighted in green. Cases where the binding pocket was incorrectly predicted are highlighted in red.

| # | Ligand | Receptor | Experimental pocket | Predicted by Boltz-2 pocket | RMSD, Å |
| --- | --- | --- | --- | --- | --- |
| 1 | WWW | TAS2R4 | extracellular | extracellular | 7.2 ± 1.4 |
| 2 | cholesterol | TAS2R14 | extracellular | - | 18.0 ± 1.0 |
| 3 | tangeretin | TAS2R14 | extracellular | extracellular | 5.9 ± 2.2 |
| 4 | chlorhexidine | TAS2R14 | extracellular | extracellular | 10.2 ± 2.4 |
| 5 | compound 28.1 | TAS2R14 | extracellular and intracellular | intracellular | 2.8 ± 0.4 |
| 6 | flufenamic acid | TAS2R14 | extracellular and intracellular | intracellular | 4.9 ± 1.8 |
| 7 | aristolochic acid | TAS2R14 | intracellular | intracellular | 6.1 ± 0.9 |
| 8 | cholesteryl hemisuccinate | TAS2R14 | extracellular | - | - |
| 9 | diiodosalicylic acid | TAS2R14 | intracellular | intracellular | - |
| 10 | salicin | TAS2R16 | extracellular | extracellular | 1.3 ± 0.2 |
| 11 | aristolochic acid | TAS2R43 | extracellular | extracellular | 3.7 ± 2.1 |
| 12 | tangeretin | TAS2R46 | extracellular | extracellular | 6.5 ± 2.4 |
| 13 | strychnine | TAS2R46 | extracellular | extracellular | 2.0 ± 0.4 |
| 14 | denatonium | TAS2R46 | extracellular | extracellular | 6.3 ± 0.1 |
| 15 | artemisinin | TAS2R46 | extracellular | extracellular | 4.1 ± 0.0 |

**Supplementary Table 2.**
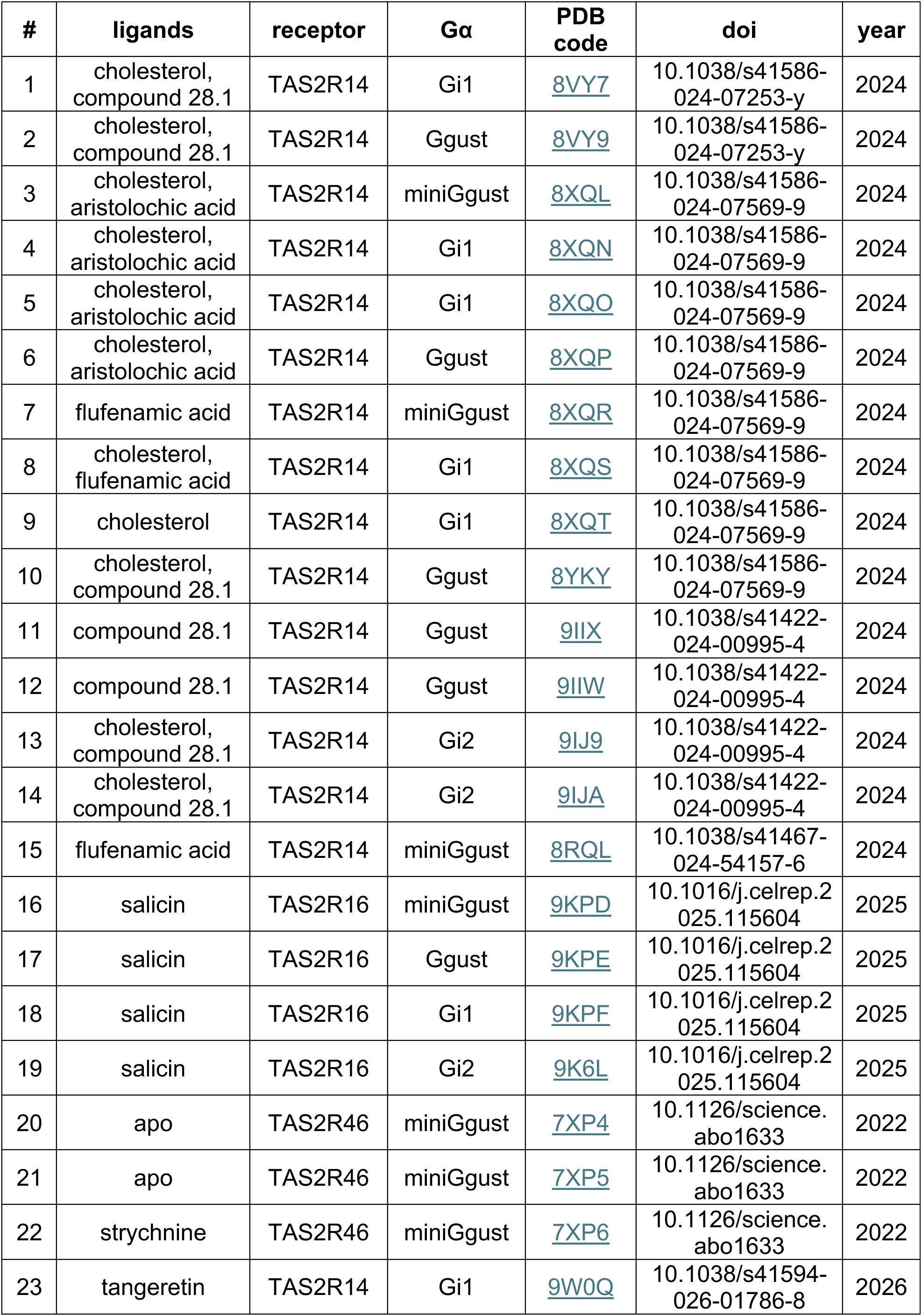
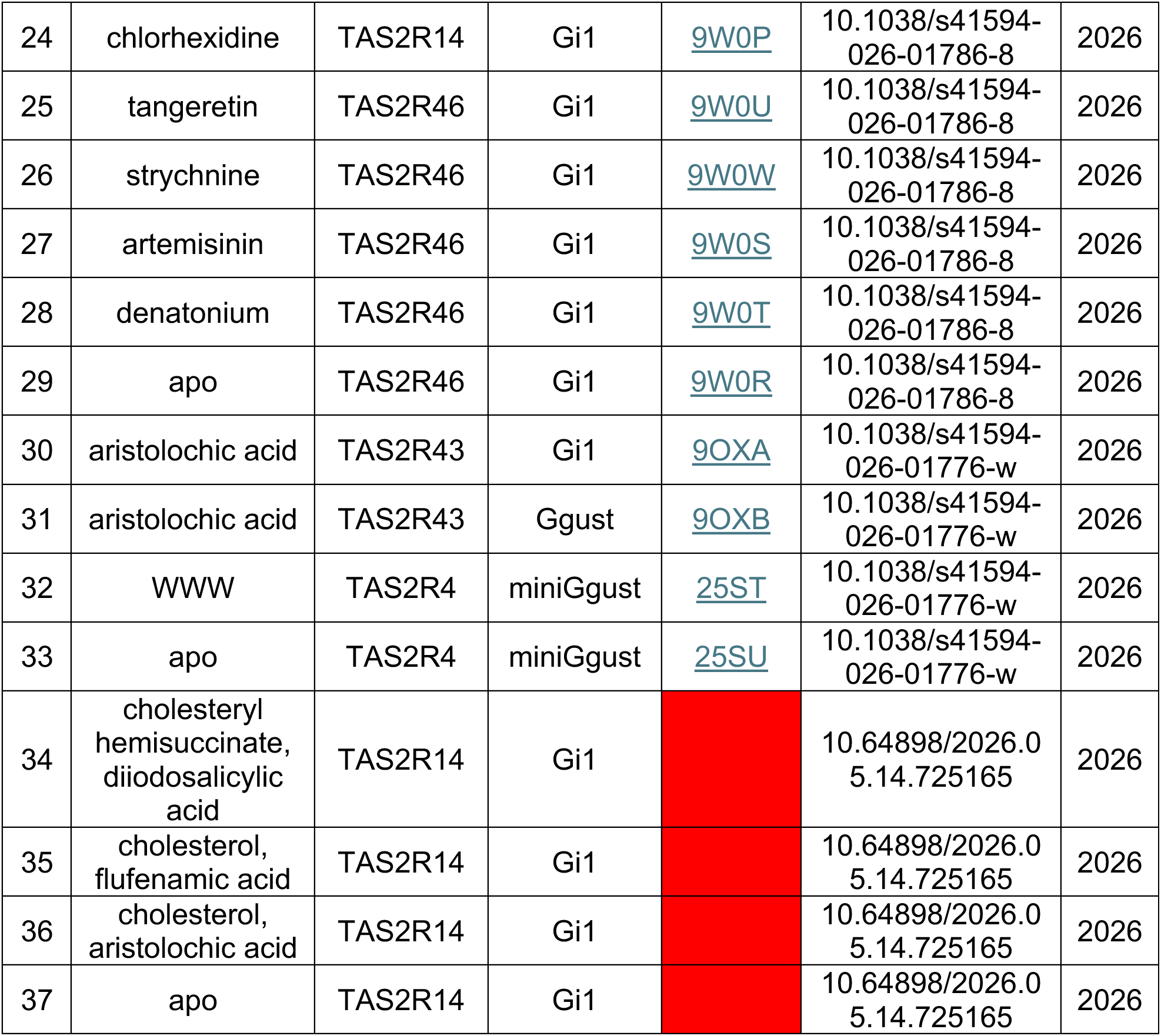
Available cryo-EM protein-ligand complexes of TAS2Rs.

**Supplementary Figure 2.**
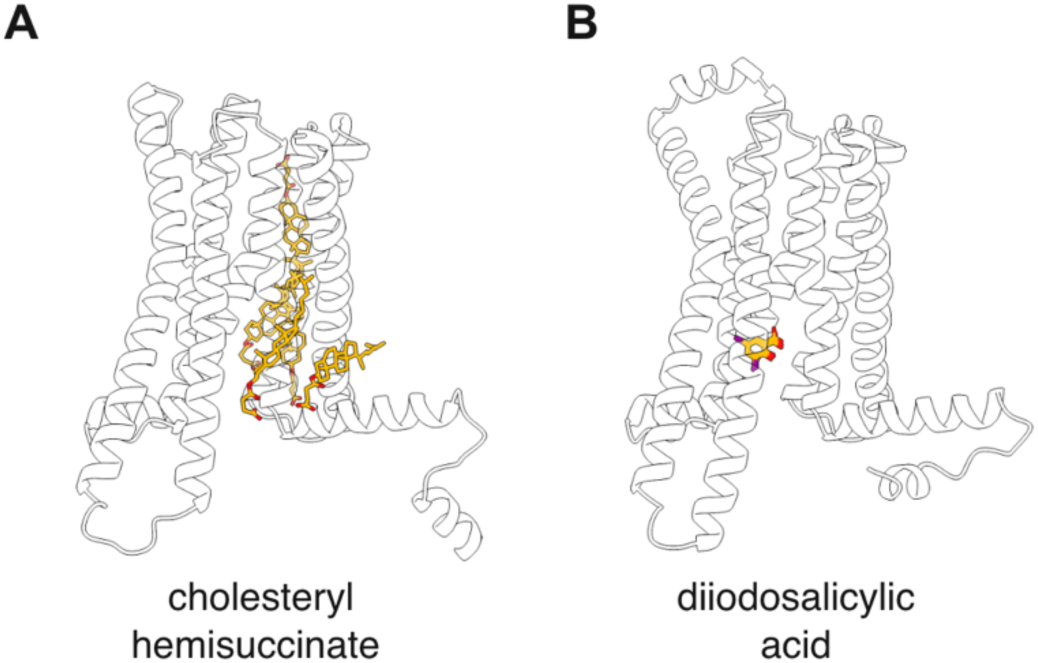
Boltz-2 predicted binding poses for TAS2R14 with (A) cholesteryl hemisuccinate and (B) diiodosalicylic acid.

**Supplementary Figure 3.**
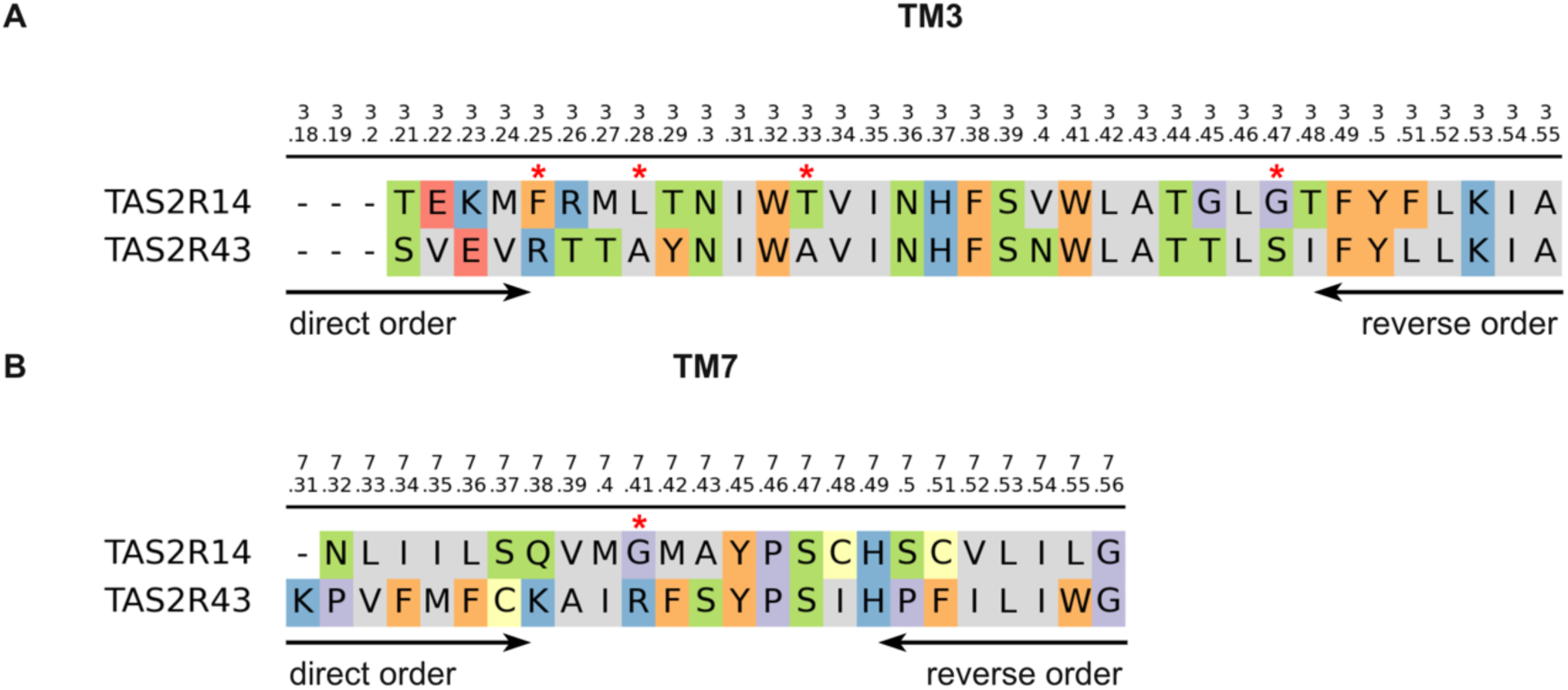
Aligned sequences of TM3 and TM7 of TAS2R14 and TAS2R43 used in step-by-step morphing. Red stars indicate the TAS2R14 residues whose mutation led to a significant increase in aristolochic acid binding in the extracellular pocket.

**Supplementary Figure 4.**
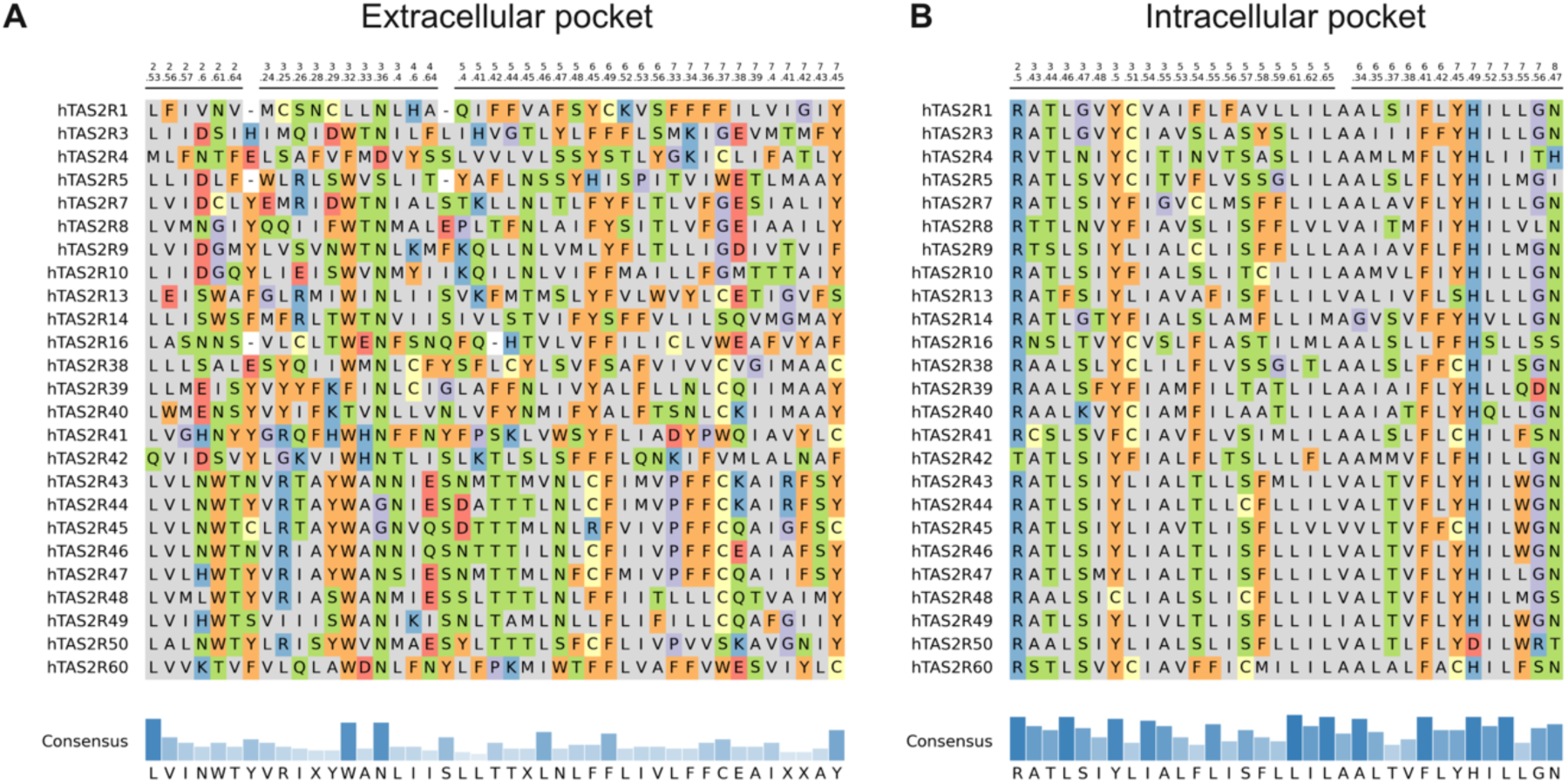
Aligned residues in the (A) extracellular and (B) intracellular pockets of human TAS2Rs.

**Supplementary Figure 5.**
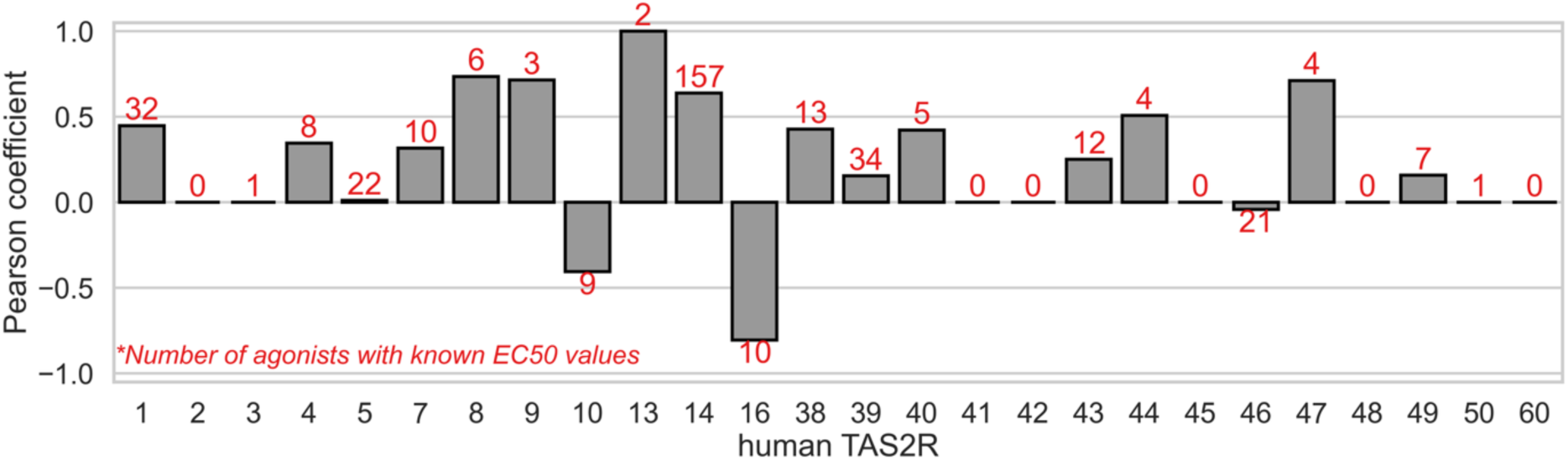
Distribution of Pearson coefficients for predicted affinity values of human TAS2Rs. The number of unique agonist-TAS2R pairs used to calculate the Pearson coefficient is shown in red.

**Supplementary Figure 6.**
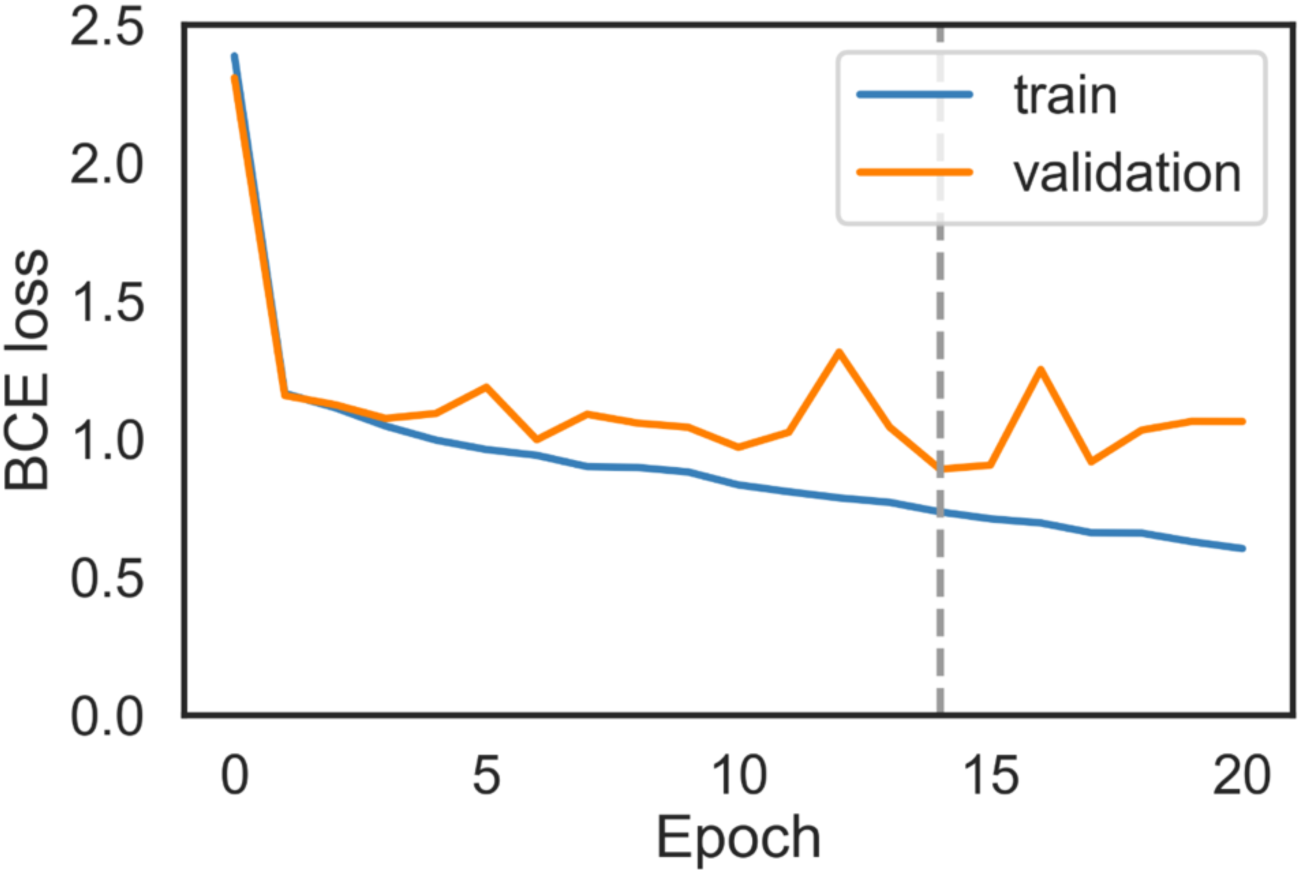
Learning curves for fine-tuning the affinity module over 20 epochs. The gray dashed line indicates the 14th epoch, which has the lowest BCE loss on the validation set.

**Supplementary Table 3.** Metrics for the best fine-tuned affinity module.

| stage | precision | recall | balanced accuracy | f1-score | roc auc | average precision |
| --- | --- | --- | --- | --- | --- | --- |
| train | 0.67 | 0.51 | 0.73 | 0.58 | 0.87 | 0.66 |
| validation | 0.6 | 0.53 | 0.72 | 0.56 | 0.82 | 0.58 |

**Supplementary Figure 7.**
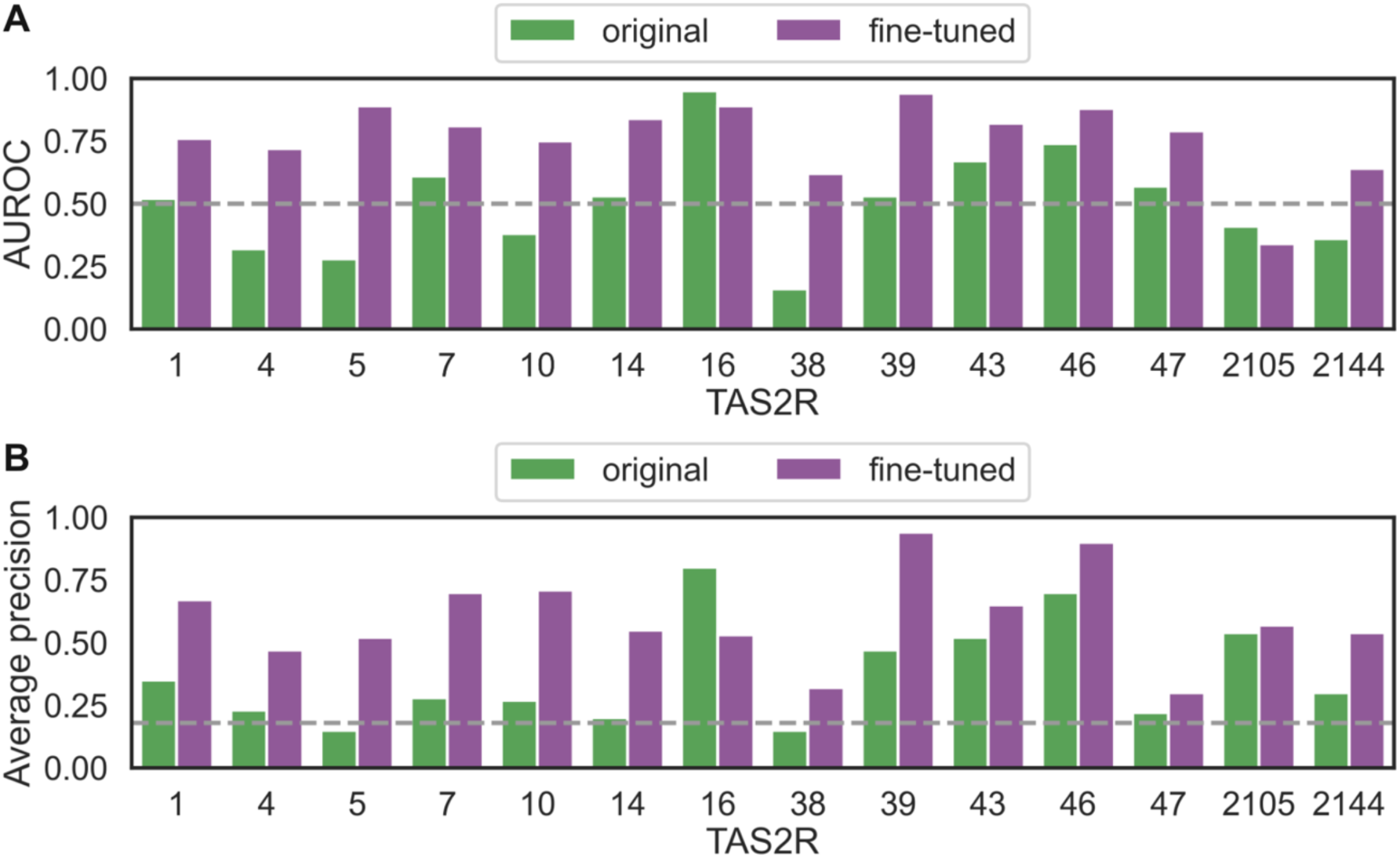
Distributions of (A) AUROC and (B) average precision for the fine-tuned affinity module for those TAS2Rs that had more than 5 agonists and 5 non-agonists in the validation set. The gray dashed line indicates the level of the random model.

